# Direct optical nanoscopy unveils signatures of cytokine-induced β-cell structural and functional stress

**DOI:** 10.1101/2023.02.20.529190

**Authors:** Licia Anna Pugliese, Valentina De Lorenzi, Mario Bernardi, Samuele Ghignoli, Marta Tesi, Piero Marchetti, Francesco Cardarelli, Luca Pesce

**Affiliations:** NEST Laboratory - Scuola Normale Superiore, Piazza San Silvestro 12, Pisa, Italy; Department of Clinical and Experimental Medicine, Islet Cell Laboratory, University of Pisa, Pisa, Italy

**Keywords:** cytokine, expansion microscopy, super resolution, metabolic imaging, β-cells, fluorescence, mitochondria, insulin secretory granules, microtubules, actin, NADH

## Abstract

Here we exploit a combination of advanced optical-microscopy tools and fluorescently-labeled molecular targets in rat Insulinoma 1E β-cells exposed to proinflammatory cytokines. Expansion microscopy (ExM) is used to achieve the spatial resolution (~50 nm) needed to analyze the structural features of key subcellular targets, i.e. insulin secretory granules (ISGs), microtubules, actin filaments, and mitochondria; time-lapse live-cell microscopy, on the other hand, provides complementary information on key dynamic and metabolic subcellular parameters. It is found that 24-hours exposure to proinflammatory cytokines induces a neat decrease in the number of ISGs and alteration in the dynamics of the residual pool, marked depolymerization of microtubules, change in mitochondrial morphology and metabolic activity, and decreased cell responsiveness to glucose stimulation. This is accompanied by clear signatures of the production of reactive oxygen species. Reported results provide direct evidence that proinflammatory cytokines act as potent stimulators of insulin secretion and, concomitantly, as cell stressors.

## Introduction

Type 1 Diabetes (T1D) is a chronic disease characterized, among others, by inflammation and consequent dysfunction of pancreatic β-cells^1^. Despite the efforts, the molecular determinants underlying β-cell failure are not fully clarified, hindering the efforts to prevent and/or cure the disease. Thus far, β-cell failure has been investigated using a combination of genomic, transcriptomic, proteomic, and biochemical approaches. Based on such “omics” data it appears clear that proinflammatory cytokines (including IL-1β, TNF-α, and IFN-γ) play a key role in β-cell failure in T1D, inducing alterations at the molecular level (e.g. by upregulation of selected genes and/or proteins^2–4^, which then, in turn, promote β-cell failure, mainly by apoptosis^5,6^. By standard ELISA-based secretion assays, it was clearly established that exposure of pancreatic β-cells to proinflammatory cytokines impair β-cell sensitivity to glucose stimulation^7,8^. On the other hand, a combination of secretion assays, RNA sequencing and mass spectrometry allowed to associate cytokine treatment and aberrant insulin secretion with the concomitant increase in proinsulin misfolding, probably due to altered proinsulin interactions with co-chaperone partner proteins, primarily within the endoplasmic reticulum (ER)^9^. Based on the discovered proinsulin ‘interactome’, the same authors also speculate on the possible cooperation of proinflammatory cytokines with microtubule destabilization to impair insulin secretion ^10^, although the relationship between microtubule-integrity regulation and insulin secretion is still highly debated, with contrasting experimental evidences reported so far^11–13^. As already noted, together with the alteration of glucose-induced insulin secretion, proinflammatory cytokines are able to induce β-cell failure. Regarding this, while the involvement of the Endoplasmic Reticulum (ER) as central hub in mediating/integrating death-inducing signals appears clear^14–17^, accumulating evidence suggests that mitochondria are additional key players in the process^6,18–22^. For instance, Barbu and co-workers showed that cytokine-induced cell failure is preceded by disruption of mitochondrial membrane potential in rat RINm5F cells^23^; similarly, Grunnet and co-workers found that cytokines are able to induce mitochondrial stress, cytochrome c release, activation of caspase-9 and -3, and finally DNA fragmentation in human and rat islets and Insulinoma 1E (INS-1E) cells^3^.

In spite of such preliminary knowledge on the potential molecular determinants and subcellular structures involved in the response to cytokines, no report thus far validated current hypotheses by direct observation of cytokine-induced alterations in β-cells by high-resolution imaging. Worthy of mention, Transmission Electron Microscopy (TEM) was used to build a large-scale imaging database using tissues from both healthy and diabetic donors^24^, but with inherently poor molecular specificity. On the other hand, exploitation of the high molecular specificity achievable by fluorescence-based optical microscopy^25^, together with the nanoscale sensitivity of super resolution methods (SRM), is still in its infancy in the study of β-cells (for a review see ref^26^). So far, in fact, fluorescence-based SRM was primarily used to investigate ISG trafficking in β-cells (ref) and its response to glucose stimulation^12,13^.

In this work, we exploit a combination of high-resolution optical-microscopy tools and fluorescently-labeled molecular targets in Insulinoma 1E (INS-1E) β-cells exposed to IL-1β and IFN-γ pro-inflammatory cytokines. Four main targets were selected: *i*) the ISG as a final effector in the process of insulin secretion; *ii*) microtubules and *iii*) actin filaments as main cytoskeleton components used by ISGs for trafficking and secretion; *iv*) mitochondria as energy factory fueling the metabolic processes of the β cell, including granule movements and insulin secretion. In more detail, expansion microscopy (ExM)^27,28^ was used to achieve the spatial resolution (~50 nm, corresponding to an expansion factor, hereafter EF, of ~4.7 folds) needed to analyze the structural features of the key subcellular targets mentioned while preserving their actual spatial organization in the specimen^29,30^. Time-lapse, live-cell microscopy, on the other hand, provided complementary information on dynamic and metabolic subcellular parameters. It was found that 24-hours exposure to pro-inflammatory cytokines induces a neat decrease in the number of ISGs (with altered dynamics of the residual ISG pool), marked alteration of cytoskeleton structure (in particular, microtubules depolymerization), change in mitochondrial morphology and metabolic function, with an overall metabolic priming of cells towards an increase in oxidative phosphorylation. These effects are accompanied by the observation of clear signatures of the production of reactive oxygen species (ROS) under cytokine exposure. Overall, reported results support the hypothesis that pro-inflammatory cytokines stimulate aberrant insulin secretion and cell stress, contributing to reduced β-cell responsivity and, eventually, failure.

## Results

### Overview of the experimental procedure

Insulinoma 1E (INS-1E) cells share many characteristics with primary β-cells (e.g. glucose-sensing ability) and are therefore widely used as a β-cell model^31^. To gain insight into the effect of pro-inflammatory cytokines on INS-1E cells, we combine immunofluorescence, which provides molecular specificity, with ExM, which physically magnifies the sample and allows super-resolved images in conventional diffraction-limited microscopes. **Fig. 1A** summarizes the experimental workflow. First, INS-1E cells were treated (or not) with pro-inflammatory cytokines for 24 hours. Cells were then chemically fixed using paraformaldehyde, glutaraldehyde, or a combination of both, to crosslink neighboring proteins^32^. Next, cells were stained using conventional antibodies functionalized with fluorescent probes, optimizing the labeling protocol for each biological target to be investigated (see Methods for more details). According to ExM standard procedure, Acryloyl-X SE (AcX) molecular handle^33^ was covalently attached to virtually any of the primary amine groups of labels and biomolecules in the sample, enabling them to be anchored to the hydrogel. At this point, the specimen was soaked in a solution consisting of acrylamide, bisacrylamide, and sodium acrylate to generate a densely cross-linked gel throughout the specimen. Such a hybrid sample was finally placed in a digestion buffer (to split up the protein content), and expanded in distilled water by exploiting the highly charged nature of the polyelectrolyte backbone^28^. Four subcellular targets were analyzed: ISGs, tubulin, actin, and mitochondria (**Fig. 1B**). The final expansion factor (EF) was determined using two independent markers, i.e. Phalloidin and TOM20, which label actin filaments and mitochondrial membranes respectively, and was found to be of 4.7±0.4 folds (**Fig. 1C**), a value that corresponds in our system to a new nominal resolution for imaging of 50±7.9 nm (**Fig. S1**). In addition, the pre- and post-expansion confocal images of the same stained cells were compared, showing low distortion and a high structural similarity index (calculated as reported in Ref^34,35^) for all the stained structures (total SSIM index = 0.73±0.05) (**Fig. S2**). With the new gained resolution, we tested confocal imaging on the four target structures selected. For what concerns ISGs, as shown in **Fig. 1D-E**, the de-crowding effect imposed by ExM favored identification of single granules and, as a consequence, their proper counting (e.g. the number of ISGs detected markedly increases in post-expansion specimens with respect to pre-expansion ones; n=25 cells, 3 independent samples). Worthy of mention, post-expansion ISGs reveal a donut-shaped staining pattern, which is not detectable in pre-expansion ones (exemplary image in **Fig. S3**): the hole of the donut might correspond to the crystalline inner core of the granule, eventually not reached by primary and secondary antibodies due to steric hindrance. For what concerns the cytoskeleton, α-tubulin is used here as a benchmark. As reported in **Fig. 1F-G**, ExM allowed to resolve individual microtubules that could not be distinguished with standard confocal microscopy (see profiles of intensity in **Fig. 1G**). Finally, as shown in **Fig. 1H**, mitochondria form a highly interconnected and dense tubular network in INS-1E cells, making it difficult to investigate their number and structural features using a conventional microscope. Analogously to previous cases, ExM allowed accurate segmentation and quantification of the number and structure of such subcellular organelles. As expected, expanded TOM20-labeled mitochondria were smaller in size (area expressed as µm^2^; **Fig. 1I**) and greater in number (**Fig. S4A**) per cell after rescaling for the EF. Also, we observed a reduced cytoplasmic area (**Fig. S4B**), indicating better individually resolved mitochondria.

**Fig. 1.**
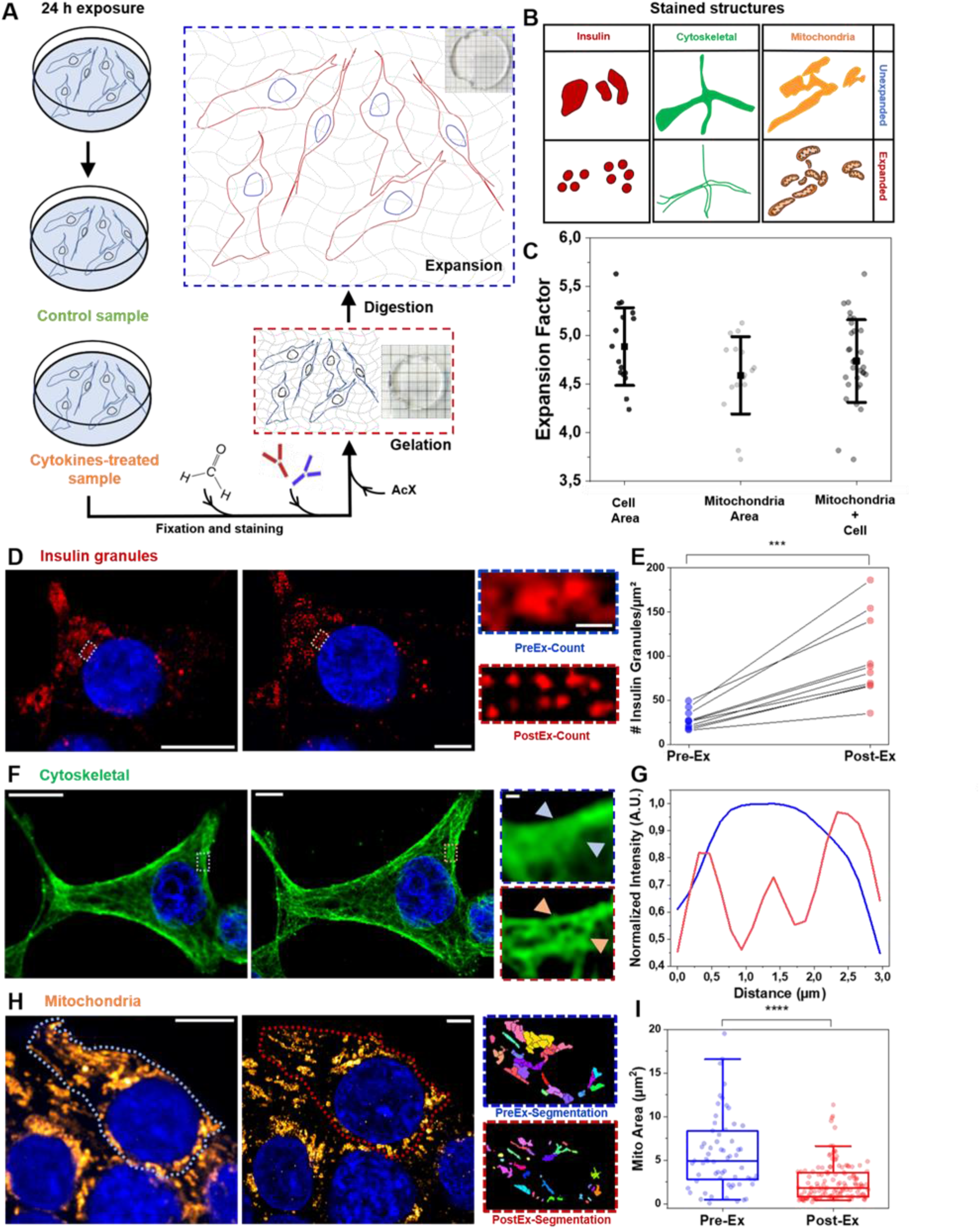
Schematic representation of the general workflow of our experiments. **A**. INS-1E were plated and then incubated for 24 h in fresh complete medium or supplemented with cytokines. Then, the specimens were chemically fixed, stained with fluorescence probes and antibodies, functionalized with AcX, gelled, and finally expanded. **B**. Cytokine-treated and control samples were stained for three subcellular structures (mitochondria, ISGs, and cytoskeletal) involved in glucose response and insulin granule tracking, and investigated with nanoscale resolution using ExM. **C**. The EF characterization was performed using subcellular (mitochondria) and cellular (actin staining) in order to achieve a more accurate EF (mitochondria + cell area EF: 4.7±0.43 SD). **D**. Pre- and post-expanded INS-1E cells labeled for insulin. The inset shows a zoomed-in region highlighting the improved resolution of expanded samples. **E**. ISG counts before versus after expansion for 25 different cells (each circle represents the single cell; n=3 independent experiments). **F**. Pre- and post-expansion confocal images of INS-1E stained for α-tubulin, with magnified views of boxed regions and (**G**) profiles of intensity taken along the blue and red arrows. **H**. Mitochondrial network confocal images and algorithm segmentation in unexpanded and expanded INS1E cells stained for TOM20. (**I**) Box plot shows morphometric analysis performed by using the MorphoLibJ plugin. The mitochondrial area is expressed as µm^2^. Data are presented as box plots with whiskers at the 5th and 95th percentiles, the central line at the 50th percentile, and the ends of the box at the 25th and 75th percentiles (n. of mitochondria = 100 for control and treated samples; 2 independent experiments). A Mann– Whitney test was performed (***P < 0.001, ****P < 0.0001). Cells were acquired by confocal microscope using 405 and 488 excitation light, with 63x/NA1.4 objective lens. Scale bar 10 µm.

### Proinflammatory cytokines promote granule secretion in INS-1E cells

It is well known that even short-term exposure to proinflammatory cytokines is able to reshape proinsulin interactions with critical regulators of the secretory pathway in β-cells, leading to hormone aberrant secretion^9,36–38^. Based on this, we started by examining the effect of cytokines on the number and localization of ISGs in cultured INS-1E cells. In detail, cells were exposed for 24 hours to cytokines (IL-1β 10 U/ml and IFN-γ 100 U/ml diluted in RPMI) in the maintenance medium (see Materials and Methods), then fixed, labeled with anti-insulin antibody, and expanded using ExM protocol. Already at a first visual inspection (exemplary images are shown in **Fig. 2A**), it appears clear that ISGs are more uniformly distributed in control cells as compared to cytokine-treated ones, in the latter being preferentially accumulated at the cell border (i.e. close to the plasma membrane). To better highlight this difference, **Fig. 2B** reports on the average fluorescence intensity recorded in a ROI drawn across the cytoplasm and borders of the cell (dashed rectangular boxes in **Fig. 2A**). As expected, in a control cell ISG-specific signal is present almost throughout the cell environment (green trace in **Fig. 2B**), while in cytokine-treated cells it is markedly higher at the borders as compared to the cytoplasm (orange trace in **Fig. 2B**). As mentioned earlier, the de-crowding effect of ExM allows a more accurate estimate of granule number in cells: to this end, the entire cellular volume was probed in expanded specimens by means of z-stack acquisitions with a z-stack step of 1 µm. Of note, a ~40% reduction of the ISGs density (i.e. ISG number/µm^2^) was observed in cytokine-exposed cells as compared to control cells (**Fig. 2C**). Such reduction in ISG content upon exposure to cytokines was confirmed by independent Western Blot analysis of the total insulin content in cell lysates (**Fig. 2D**) and supports the current model of aberrant insulin secretion induced by cytokines^9,36–38^. To complement the information on ISG density and localization with dynamic information on the population of residual granules, we performed live-cell time-lapse acquisition and iMSD (i.e. imaging-derived Mean Square Displacement^39,40^), analysis of granule motion. **Fig. 2E-G** summarizes the workflow of the iMSD experiment: ZIGIR^41^, a fluorescent granule indicator with Zn^2+^-chelating properties (see Materials and Methods) (**Fig. 2E**) is used to fluorescently label ISGs in live cells; a stack of images is acquired at ~200 ms/frame temporal resolution for at least 100 seconds in a 12×12-µm^2^ region of interest (**Fig. 2F**); the spatiotemporal correlation function is calculated and interpolated by a Gaussian function; the iMSD trace is finally obtained by extracting the variance of the Gaussian fit at each time delay (**Fig. 2G**). Thanks to the iMSD analysis, the average dynamic (i.e. anomalous diffusion parameter α and local-diffusivity coefficient Dm) and structural (i.e. organelle size through the iMSD y-axis intercept) were extracted and used to compare the cytokine-exposed and control conditions. While granule average size is not affected by cytokines (**Fig. S5**), both the dynamic parameters show statistically significant alterations (**Fig. 2F-G**): in particular, the α coefficient decreases under cytokine treatment while, concomitantly, Dm increases. These data suggest that the residual granule population after 24-h exposure to cytokines is less prone to perform active transport as compared to granules from untreated cells

**Fig. 2.**
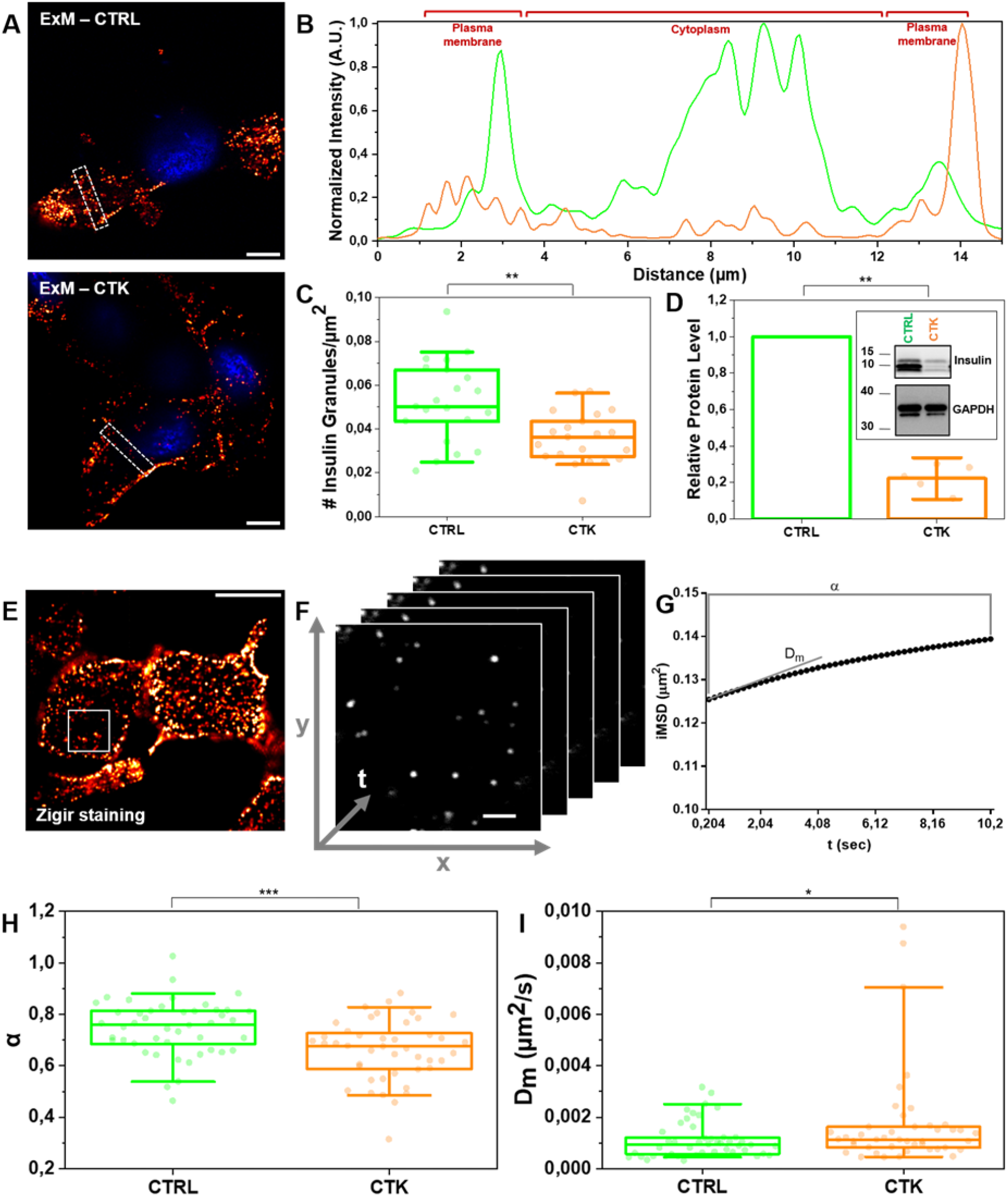
Proinflammatory cytokines induce ISGs degranulation. **A**. Control and cytokine-treated sample of INS-1E cells stained for ISGs in maintenance condition. Cells were acquired by confocal microscope using 405 and 488 excitation light, with 63x/NA1.4 objective lens. Scale bar 10 µm. **B**. Profiles of intensity taken along the 2×15 µm^2^ white square. The plot profiles show a more homogeneous distribution of IGs in the cytoplasm in the control (green) with respect to cytokine-treated samples (red). **C**. Granule count in CTRL and CTK expanded samples (ExM), showing a statistically significant reduction of the insulin content in CTK samples. Data were presented as box plots with whiskers at the 5th and 95th percentiles, the central line at the 50th percentile, and the ends of the box at the 25th and 75th percentiles (n=21 cells; 3 independent experiment). **D**. Total proteins were extracted, and the expression of insulin were assessed by Western blotting (WB). GAPDH was used as a control for protein loading. Protein signals were quantified and corrected for the corresponding GAPDH value and expressed as fold change compared to untreated cells (CTRL) (n = 5 independent experiments). (**p ≤ 0.01; Non-parametric Mann Whitney test; significantly different from the control condition at 24 h of incubation). iMSD explanatory figure: **E**. Cells marked by ZIGIR fluorophore were imaged (2x image scale bar 10 μm) with 500 frames time-lapse images **F**. Example of a stack of images acquired at 204.80 ms per frame. Scale bar 2 μm. **G**. iMSD analysis pointed out a granule motion descriptor (α) and diffusivity coefficient (Dm) for each time-lapse. **H**. Comparison of the α parameter between control (CTRL, n = 46) and cytokines treatment (CTK, n = 44) (3 independent experiments). **I**. Comparison for the diffusion coefficient (Dm) between CTRL (n = 46) and CTK (n = 44) (3 independent experiments). We performed the Shapiro-Wilk normality test, unpaired t test for α and Kolmogorov-Smirnov test for Dm based on the normality (see Materials and Methods).

### Proinflammatory cytokines promote cytoskeleton rearrangements: microtubules and actin filaments

The cytoskeleton is an essential regulator of ISG intracellular trafficking and localization. Several studies demonstrated that microtubules density can affect glucose-stimulated insulin secretion (GSIS): in fact, a reduction in microtubules density increases GSIS while, by contrast, a high microtubules density inhibits GSIS^11,13,42,43^. Similarly, Tran and collaborators observed that chemical destabilization of microtubules by Nocodazole is associated with an increased proinsulin secretion in response to cytokines^9^. These reports prompt us to speculate that the reduced number and altered dynamics of granules under cytokine exposure observed here may be correlated with changes in microtubules structure. To analyze the impact of cytokines on MT organization, we selected three different 10×10 µm^2^ Regions Of Interest (ROIs) within a complete Z-stack acquisition of each cell. These ROIs were: i) basal tubulin – No Nuclear Localization (NNL) (α-tubulin found in contact with the glass and in the first planes of the Z-stack), ii) basal tubulin – Nuclear Tubulin (NT) (α-tubulin found in proximity of the glass and beneath nuclei), and iii) equatorial tubulin (ET), both in normal and cytokine-treated samples (shown in **Fig. 3A** for a representative cytokine-treated INS-1E cell). For each cell, we extracted more than one ROI from the basal and equatorial cytoplasm to ensure that we covered the entire cell surface and obtained an optimal signal-to-noise ratio for efficient tracing (**Fig. 3A**). The expanded images in **Fig. 3B** for INS-1E cells labeled for α-tubulin show a significant decrease in MT density in cells treated with cytokines (bottom panel) compared to control cells (upper panel). To quantify these changes, we used the FiNTA algorithm to trace the cytoskeletal network in 10×10 µm^2^ ROIs (**Fig. 3C**)^44^. This generated a binary mask of the network (**Fig. 3D**) that we further processed resorting to the Skeletonize function^45^ to extract information about an ensemble of parameters that describe the structure of the cytoskeleton, including the number of branches (portion of segments connecting by end-points, or endpoints – junctions, or junctions – junctions; brown in **Fig. 3D**), junctions (green), triple points (yellow) and quadruple points (red) (junctions with exactly 3 and 4 branches, respectively), endpoints (in blue), as well as the average lengths of branches (**Fig. 3E**). Exposure to cytokines significantly reduced the number of branches (control: 198.0 ± 4.7, cytokines: 125.0 ± 5.3; reduction of 37%) (**Fig. 3F**), junctions (control: 115.1±2.8, cytokines: 69.3 ± 3.3; reduction of 40%) (**Fig. 3G**), triple points (control: 108.1 ± 2.7, cytokines: 65.8 ± 3.0; reduction of 39%) (**Fig. 3H**), quadruple points (control: 6.6 ± 0.3, cytokines: 3.4 ± 0.3; reduction of 48%) (**Fig. 3I**), endpoints (control: 43.3 ± 1.2, cytokines: 38.6 ± 1.1; reduction of 11%) (**Fig. 3J**). Furthermore, we observed a significant increase in the average branch length (control: 0.330 ± 0.004 µm, cytokines: 0.380 ± 0.007 µm; reduction of control 13%) (**Fig. 3K**). Finally, binary images were processed with MorphoLibJ (Fiji plugin, see Methods) to segment the MT network and calculate the number, area (**Fig. 4A**) and perimeter of MT meshes (**Fig. S6**). This analysis revealed an increase in the mean area number in control samples (**Fig. 4B**), with an average spacing of 0.260 ± 0.004 µm^2^ (**Fig. 4C**). In contrast, cytokine-treated samples had a decrease in mean area number (**Fig. 4B**) and a twofold increase in the MT opening size (0.487 ± 0.010 µm^2^) (**Fig. 4C**). This structural outcome was also confirmed by the intensity distribution generated through the repetition of 175 lines running across the center of the frame (**Fig. 4D** and **Fig. S7**). Statistical analysis of the number and position of peaks along these lines indicated that cytokines-treated samples had a lower number of MT filaments compared to control samples (**Fig. 4E**), with a greater distance between peaks (**Fig. 4F**). Western blot analysis of α-tubulin shows that cytokine treatment does not reduce the cytoskeletal content in INS-1E cells (**Fig. S8A**). Such a MT remodeling can be explained with depolymerization and/or spatial re-organization of the cytoskeletal mesh. By modeling the cytokine-induced MT alteration using a basic network (**Fig. S9**), we confirmed that the MT destabilization can be interpreted as a depolymerization process, which leads to dysregulate insulin secretion in absence of direct glucose stimulation.

**Fig. 3.**
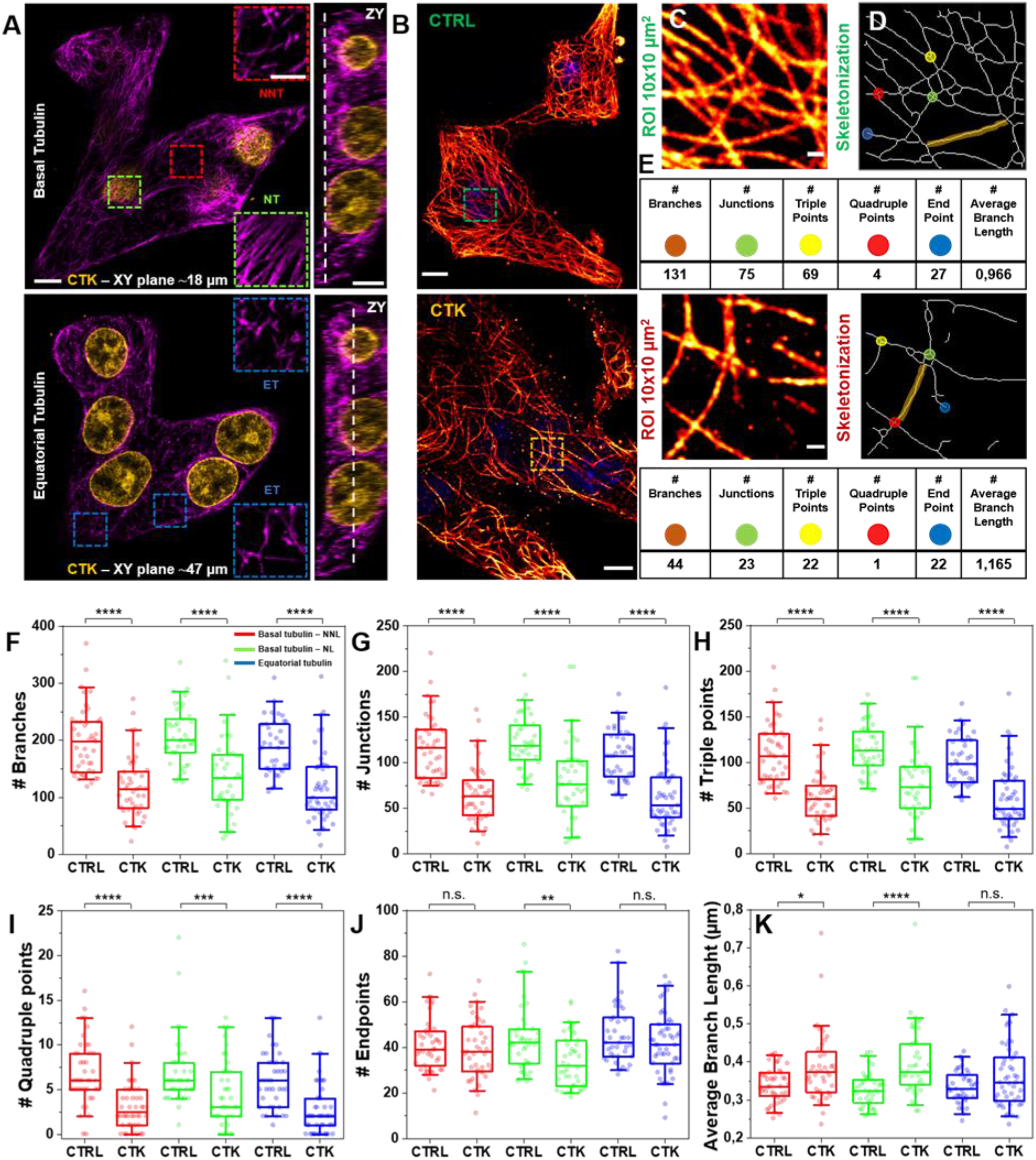
Proinflammatory cytokines promote microtubule rearrangement. **A**. Representative Z-stack image of treated sample and ROI collection at different planes. ROI were taken at the basal plane (Z-plane ~18 µm) - in the proximity of the nucleus (termed Nuclear Tubulin; NT) and outside the nucleus (termed No Nuclear Tubulin, NNT) - and at the equatorial level (Equatorial Tubulin, ET; Z-plane ~47 µm). The ZY projection shows an example of the localization along the Z-axis. Scale bar 10 um. **B**. Representative images and microtubule-network analysis of expanded INS-1E cells untreated (control, CTRL) and treated with cytokines (CTK). Cells were stained for α-tubulin and DAPI and acquired by confocal microscope using 405 and 488 excitation light, respectively, with 63x/NA1.4 objective lens. **C**. ROI of 10×10 µm^2^ were then analyzed by FiNTA and skeletonized through Fiji (**D**), to quantify the reduction in the number of branches (# Branches), junction (# Junctions), triple and quadruple points (# Triple and Quadruple points), endpoints (# Endpoints), and average bench length (**E**). Scale bar 10 µm. Quantification of MT density per ROI: # Branches (**F**), # Junctions (**G**), # Triple (**H**) and Quadruple (**I**) points, # Endpoints (**J**), and Average Branch Length (**K**). Cytokine-treated specimens were significantly different from the control condition in each plane. In addition, the average branch length increases in cytokine treated samples with respect to the control. Data were presented as box plots with whiskers at the 5th and 95^th^ percentiles, the central line at the 50th percentile, and the ends of the box at the 25th and 75th percentiles (n ≥ 39; 3 independent experiment). A Mann–Whitney test was performed.

**Fig. 4.**
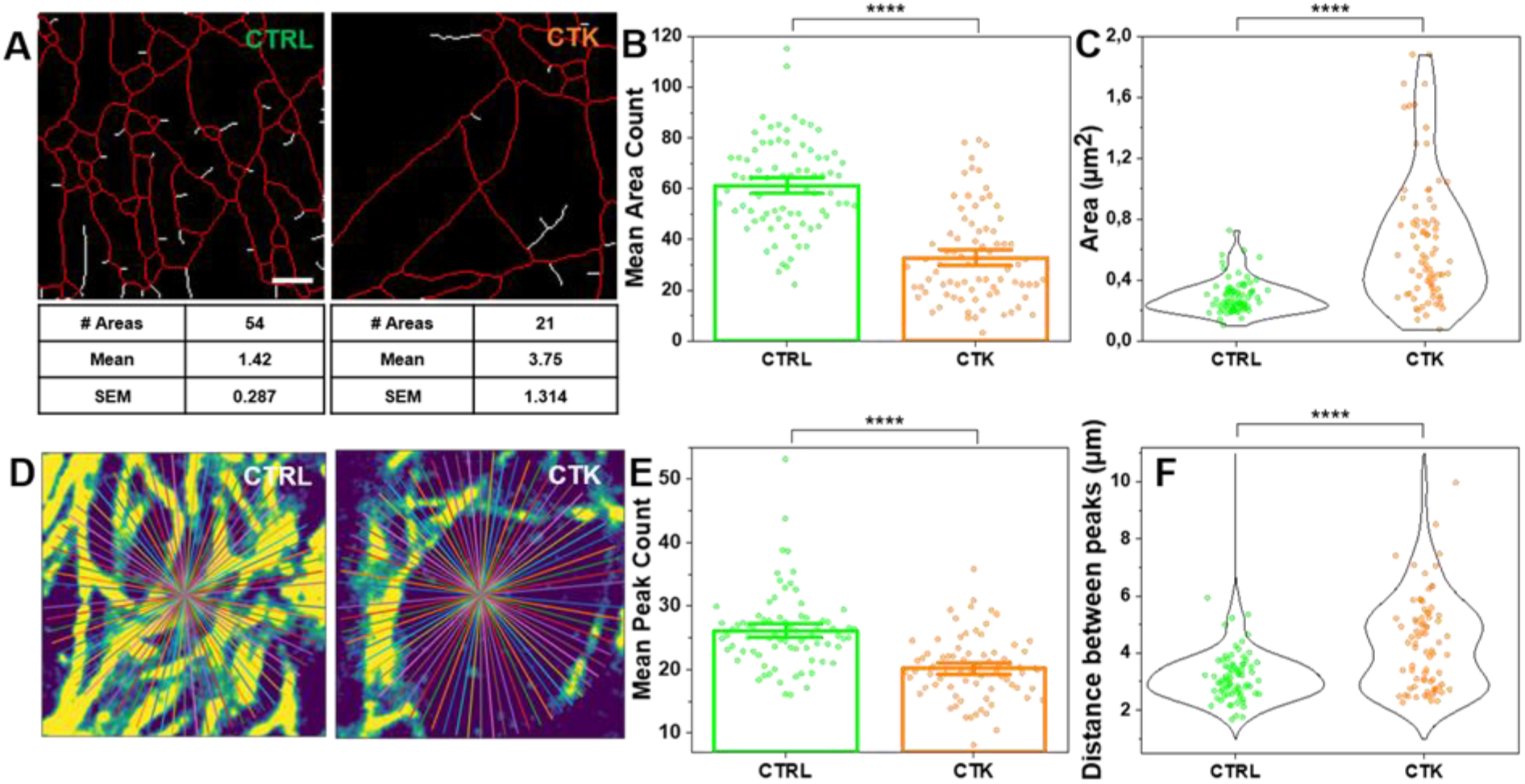
Proinflammatory cytokines promote microtubule rearrangement. **A**. MT opening analysis by using MorphoLibJ algorithm. The number of opening and mean areas were calculated for each ROI in control and treated samples. Bar ± SEM (**B**) and Violin plot (**C**) show the number of MT openings and mean area, respectively, for each ROI. (n = 76; 3 independent experiments). (**D**) Intensity distribution analysis of 175 lines passing from the frame center in control and cytokine-treated samples. Bar ± SEM (**E**) and violin plot (**F**) show the mean peak count and mean distance between peaks per ROI. A Mann–Whitney test was performed. Scale bar 10 µm.

Cortical actin is typically defined as a thin layer of dense actin filaments in close proximity (approximately 10-20 nm) to the plasma membrane of the cell^46,47^ and forming a mesh of small fenced regions (termed corrals) within which membrane and/or cytoplasmic proteins/organelles can become transiently trapped^48^. Phalloidin, typically used to label cortical actin in cells, lacks amine groups for gel anchoring: to tackle this issue and retain the fluorescence signal after expansion, in the first step fluorophore-conjugated phalloidin was labeled using anti-fluorophore antibodies, followed by hydrogel synthesis, enzymatic digestion, and expansion (see Materials and Methods for more details). After confocal acquisition, the images were processed by Finta, and skeletonized through Fiji, in order to calculate filament length, number of branches, junctions, triple and quadruple points, endpoints, shape and size of actin corrals. Since cortical actin shows a diverse and heterogeneous arrangement in INS-1E cells (**Fig. 5A**), the entire cell was analyzed and results then normalized by the cell area (and by the EF). Already at a first visual inspection, contrary to microtubules, the cortical-actin organization did not show significant alterations induced by cytokine treatment (**Fig. 5A**). This was confirmed quantitatively by Finta skeletonization and MorphoLibJ analysis (Fiji plugin, used to segment corrals and calculate the area and number, see Materials and Methods for more details): both tools yielded no significant modifications of the selected actin structural parameters (results are reported in **Fig. 5B - E**). These results from imaging are accompanied by WB analysis which detected no variation in the total amount of actin (**Fig. S8B**). Although the nanoscale intracellular organization of the actin meshwork is not affected by cytokines, a substantial increase in the number of micrometric cytoplasmic protrusions was observed in treated cells as compared to control ones (**Fig. 5F**). Such structures, also known as cytoplasmic “blebs”, are a special type of cell protrusions that are driven by intracellular pressure or caused by oxidative-stress^49^, and typically enriched in actin filaments^50^. When INS-1E cells were treated with cytokine for 24 h, a massive increase in the incidence of surface blebbing was observed (~72%), (**Fig. 5F** and **5G**).

**Fig. 5.**
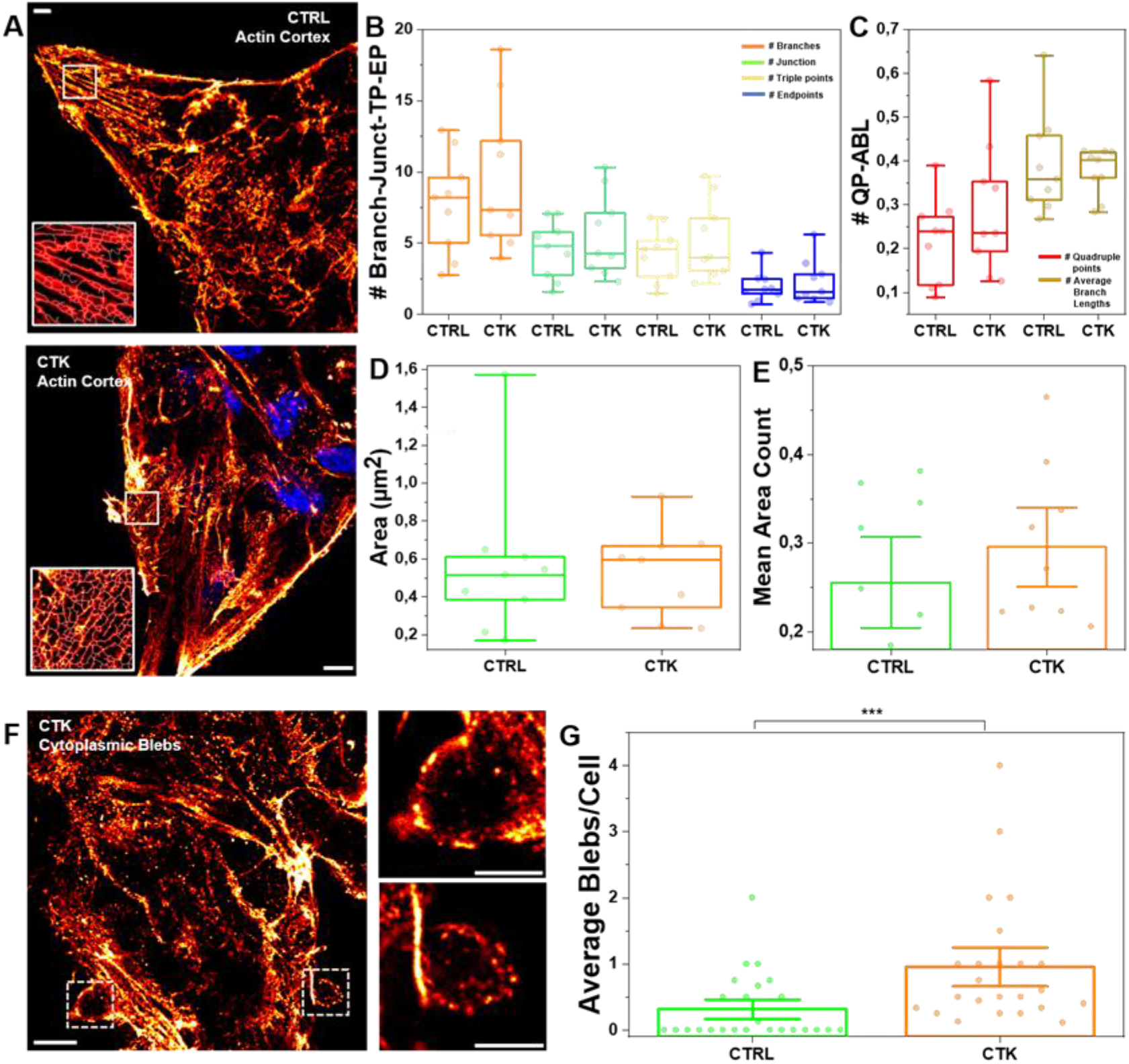
Super-resolution images of cortical actin do not show significant structural alteration. **A**. Representative images of the actin meshwork in expanded INS-1E cells untreated (control, CTRL) or treated with cytokines (CTK). Cells were stained for Phalloidin and DAPI and acquired by confocal microscope using 405 and 488 excitation light, respectively, with 63x/NA1.4 objective lens. The entire cell clusters were then analyzed by FiNTA and skeletonized through Fiji, to quantify the number of branches (Branch), junction (Junct), triple points (TP), endpoints (EP) (**B**), quadruple points (QP) and average length branch (ALB) (**C**). No significant differences in the cortical actin density were highlighted. Data were presented as box plots with whiskers at the 5th and 95th percentiles, the central line at the 50th percentile, and the ends of the box at the 25th and 75th percentiles (n=28; 3 independent experiment). Scale bar 10 µm. (**D**) Cortical actin opening (corrals) analysis and (**E**) corrals count performed by using MorphoLibJ algorithm. Data were presented as box plots with whiskers at the 5th and 95th percentiles, the central line at the 50th percentile, and the ends of the box at the 25th and 75th percentiles (n=28) (**D**); Bar ± SEM (M) shows the number of corrals. **F**. Representative image showing pathological blebs in cytokines treated sample and bleb count analysis (**G**), shown as bar ± SD (n=93). A Mann–Whitney test was performed (***P < 0.001).

### Proinflammatory cytokines induce mitochondrial structural and functional alterations: ExM and phasor-FLIM signatures

Several biochemical processes are integrated within mitochondria, from the activation of pro-apoptotic pathways to producing important metabolites, including ATP, via cell respiration. In addition, an increasing number of studies place mitochondrial dysfunction as an hallmark of Diabetes pathogenesis^18–20^. After 24 hours of cytokine treatment, cells were fixed, immunostained by TOM20, a component of the translocase of the outer membrane (TOM) complex, and, finally, expanded. Then, cytokine-treated cells were compared to the control ones for mitochondria morphometric investigation. As shown in **Fig. 6A**, the classical filamentous pattern of labeled mitochondria appears fragmented into numerous small punctate particles upon exposure to cytokines. This evidence mirrors quantitatively into the analysis of mitochondrial area and circularity parameters (**Fig. 6B-C**): the former changes from 2.88±0.29 µm^2^ in control cells to 1.29±0.07 µm^2^ in treated cells (reduction of 55%), the latter from 0.43±0.03 in control cells to 0.77±0.02 in treated cells. Overall, mitochondria in treated cells appear smaller in size and altered in shape, structural features typically associated with dysfunctional mitochondria^51^ and, presumably, metabolic stress. To probe this possibility, we related ExM-derived structural parameters to a functional assessment of cell metabolic activity performed by Fluorescence Lifetime Imaging Microscopy (FLIM) microscopy on the intrinsic signal of NAD(P)H molecules in live cells^52,53^. In fact, FLIM on NAD(P)H signal is a label-free approach used to interrogate cell metabolism which exploits the discrimination of the free and protein-bound forms of NAD(P)H molecules based on their difference in auto-fluorescence lifetime^52,53^. In previous works by some of us, the method was used to test β-cell responses to glucose stimulation both in INS-1E cells^40^ and in human-derived islets^54^. Here NAD(P)H lifetime imaging was performed in maintenance conditions (glucose concentration of 11 mM), and under pulsed glucose stimulation (shift in glucose concentration from 2.2 mM to 16.7 mM), both in control and cytokine-treated cells. NAD(P)H species were selectively excited at 740 nm and fluorescence collected at 420-460 nm; finally, the well-established, fit-free, and graphical phasor-based approach to FLIM was used to analyze data^55^. To start, we compared the metabolic state of control cells in maintenance culturing conditions (11-mM glucose concentration in RPMI medium) to that of cytokine-treated cells under the same culturing conditions (**Fig. 6D**). Interestingly, cytokine-treated cells show a neat shift towards the bound form of NAD(P)H with respect to control samples (**Fig. 6E**): this is highlighted in **Fig. 6D** by means of a color-coded map of intracellular NAD(P)H bound/free ratio. Also, cumulative results from the whole population of acquired cells (n=16 cells from n=3 independent experiments) are reported in **Fig. 6F**. Once assessed such a ‘priming’ of cytokine-treated cells towards bound NAD(P)H, we tested cell response to glucose pulsed stimulation (see Materials and Methods for details on the protocol applied). In this experiment, cytokine-treated cells show a markedly reduced metabolic response to glucose in terms of phasor shift toward higher values of the bound/free NAD(P)H lifetime ratio (**Fig. 6G**). Interestingly, such a reduced response to glucose correlates with intracellular oxidative stress (**Fig. 6H**). Indeed, the cumulative phasor plot of the whole cytokine-treated sample (n=16; frames from 3 independent experiments) contains a mixture of three species (**Fig. 6L**) respect with control (**Fig. 6I**), namely oxidized lipid-associated fluorescent species (LLS), characterized by a lifetime of approximately 8 ns^56^ (**Fig. 6I** and **S10**), free NAD(P)H, and protein-bound NAD(P)H. The ‘elongation’ of the lifetime distribution toward the LLS-specific position in the phasor plot in cells exposed for 24 h to cytokines is highlighted using a yellow cursor, while the metabolic trajectory is shown using the same color scheme represented above.

**Fig. 6.**
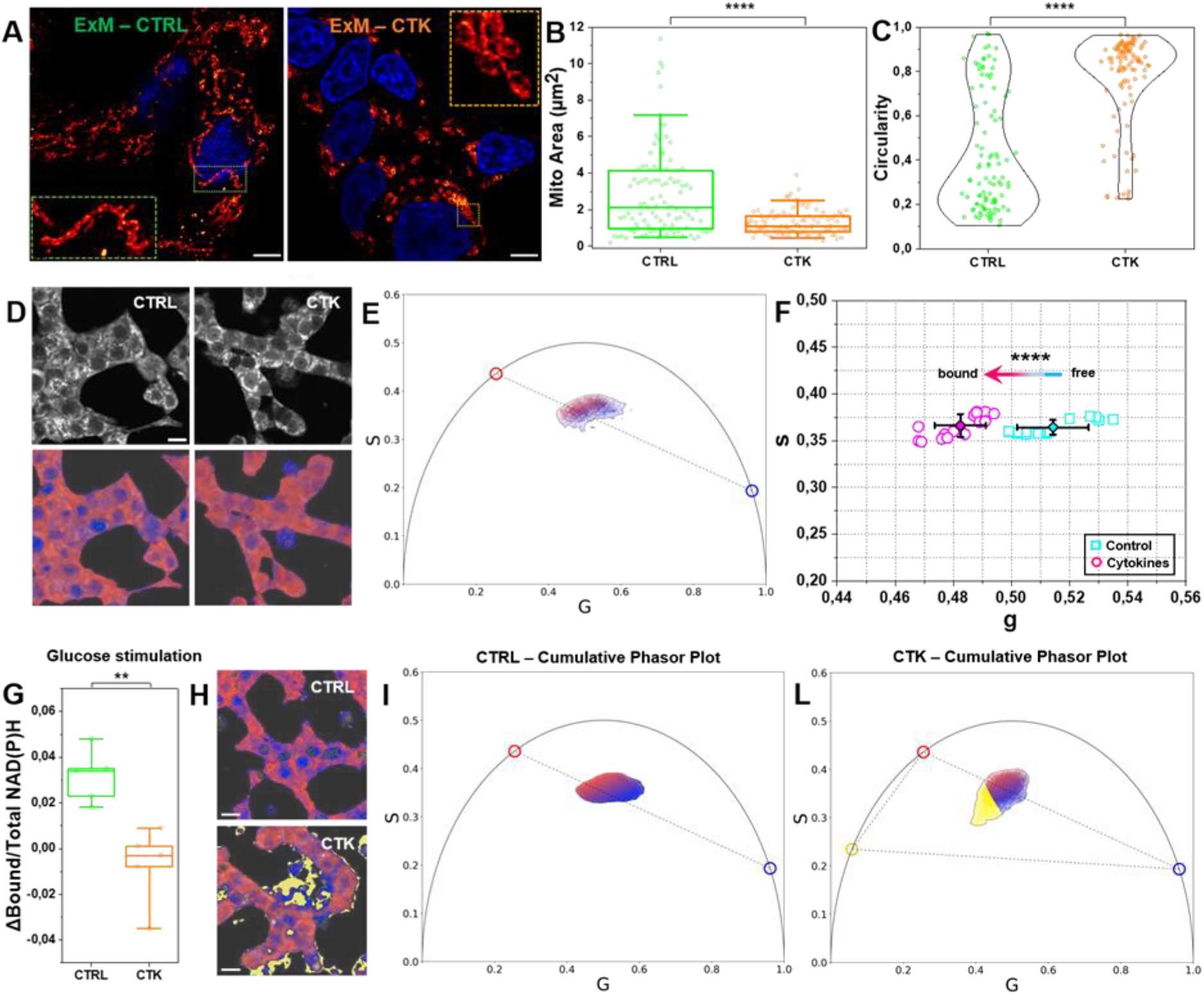
ExM and Phasor-FLIM analysis for mitochondria investigation. **A**. Representative images of expanded INS-1E cells stained for TOM20 after incubation with fresh medium (CTRL) and cytokines (CTK) for 24 h. Scale bar 10 um. The structural analysis on expanded mitochondria shows a reduced area (**B**) and high circulatory value (tending to 1) (**C**) in cytokine-treated samples. **D**. Exemplary images of total NAD(P)H intensity of live INS-1E cell clusters. As shown on the bottom line, the color-bar defines the metabolic path from NAD(P)H in the bound state (magenta) to NAD(P)H in the free state (green). **E**. Phasor plot of control and cytokine-treated cells for all pixels of acquired image in **D**. The blue and red circles represent the position in the phasor plot of pure NADH and bound NADH to lactate dehydrogenase in solution, respectively. **F**. Scatter plot of the average values of distinct phasor distributions, each relative to distinct acquired cells at the maintenance conditions. Blue squares represent control, while magenta circles show the cytokine-treated samples. Mean and SD were representative in the scatter plot (n = 16 frames; 3 independent experiments). A Mann–Whitney test was performed (****P < 0.0001). **G**. Metabolic response for glucose stimulation (n = 5; 2 independent experiments) shows a NADH bound-shift in control sample, while for cytokine-treated samples is approximately zero. Data were presented as box plots with whiskers at the 5th and 95th percentiles, the central line at the 50th percentile, and the ends of the box at the 25th and 75th percentiles. A Kolmogorov–Smirnov test was performed (**P < 0.01). **H**. LSS species as oxidative stress signatures induced by cytokines. Exemplary images and cumulative lifetime-based color map for NADH metabolic trajectories of control (**I**) and treated-samples (**L**). Pixels with LLS species are colored in yellow, according to the position of the yellow cursor in the phasor plot.

## Discussion

In this work, we exploit a combination of high-resolution optical-microscopy tools and fluorescently-labeled molecular targets in Insulinoma 1E (INS-1E) β-cells exposed to IL-1β and IFN-γ pro-inflammatory cytokines.

The starting point of our investigations was the ISG. The proposed combination of ExM-based imaging of fixed cells at 50-nm spatial resolution and iMSD-based analysis of live-cell time-lapse acquisitions allowed for an accurate estimate of ISG number, size and dynamics under cytokine-exposure and control conditions. While confirming the ability of cytokines to induce aberrant insulin secretion (here denoted by a markedly decreased number of ISGs after 24-hours exposure to cytokines), our experiments uncover altered dynamic properties in the residual population of granules: indeed they appear more dynamic (increased local diffusivity) but at the same time less prone to perform active transport (as indicated by a decrease in the anomalous coefficient alpha) compared to those exposed to control conditions. A similar change in dynamics was reported by some of us for ISGs altered by overexpression of the Phogrin transmembrane or by cholesterol overload^40^, both of which are associated with decreased cell responsiveness to glucose stimulation, in line with what observed here testing the metabolic response of cells to glucose after 24-hour exposure to cytokines. Such results on ISGs (i.e. altered secretion and altered dynamics of the residual population) prompted us to evaluate the structural properties of the cytoskeleton components, with a particular focus on microtubules. In fact, microtubules proved to act as fundamental tracks for granule mobilization, although their structural-integrity regulation during secretion remains elusive, due to the contrasting results reported so far: in fact, while Heaslip and co-workers found increased microtubule polymerization upon glucose stimulation (with negligible rearrangements of the actin mesh^12^), other reports indicate that microtubule hyper-stabilization prevents and microtubule depolymerization promotes the capacity of β cells for GSIS; interestingly, massive nocodazole-induced microtubule depolymerization (reminiscent of the cytokine effect observed here), extends the first phase of GSIS and causes oversecretion^11,13^. This latter idea of microtubule depolymerization acting as an enhancer of insulin secretion is supported by what is observed here in cytokine-exposed cells. In fact, ExM-derived structural parameters are compatible with microtubule rearrangements in terms of depolymerization (no variations of the tubulin total content were observed). Please note that microtubules depolymerization in cytokine-treated cells is also compatible with the presence of a pool of intracellular granules with decreased active transport and increased local diffusivity. The similarity between the effects of cytokines and those of glucose are not limited to what is observed for microtubules. In our experimental system, in fact, cytokine-induced insulin secretion is also associated with a cell metabolic status that recalls the one previously measured under glucose induced insulin secretion^57^. In both cases, the stimulus induces a neat increase of the ratio between protein-bound and free NAD(P)H lifetimes, classically interpreted as an increase in the oxidative-phosphorylation activity of the cell^54,57^. Based on the current model of β-cell function, the shift from low to high bound/free NAD(P)H lifetime ratios well agrees with the need to rapidly increase the ATP/ADP ratio by oxidative phosphorylation to then stimulate the cascade of biochemical reactions leading to insulin secretion. In this picture, it is not surprising that glucose and cytokines stimuli, both capable of promoting insulin secretion, share the same metabolic fingerprint; at the same, β-cell priming to oxidative-phosphorylation, besides sustaining a secretion step, inevitably makes the cell less responsive to further stimulation by glucose (as measured here and confirmed by others by independent approaches, ref). ExM-based super-resolution imaging of mitochondria highlights that their peculiar metabolic (functional) status under cytokine exposure is associated with structural alterations, i.e. fragmentation of the mitochondrial network and a neat change in mitochondrial shape. An additional hallmark of cytokine-exposed β cells emerging from previous biochemical investigations is their high degree of susceptibility to oxidative damage, primarily due to deficiency of antioxidant enzymatic machinery to counteract reactive oxygen species (ROS)^58,59^. For instance, Miki and co-workers demonstrated higher vulnerability to oxidative stress of β-cell with respect to alpha cells, because of a lower expression of GPx and catalase^60^. Label-free infrared microscopy, in addition to direct imaging of NAD(P)H lifetimes, offers the exquisite sensitivity to ROS presence and activity, in particular to ROS-generated lipid-oxidation products, by virtue of the characteristic long lifetime of these latter^56^. Interestingly, INS-1E cells treated with cytokines show a high level of this biomarker of oxidative stress.

In conclusion, from a biomedical perspective, the results reported here in a simplified model of β cells set the guidelines for data acquisition and interpretation in future studies. An exciting possibility using such methodologies would be that of studying β cells in the natural context in which they operate, i.e. the intact Langerhans islet from ex-vivo fresh samples or paraffin-embedded tissues, potentially derived also from donors naturally exposed to cytokines (e.g. T1D donors). In this sub-millimetric organ, the presence of multiple cells tightly packed in a three-dimensional architecture makes it even more complicated to achieve single-cell identification and sub-cellular analysis, mainly because of the refractive-index mismatch caused by the variety of biomolecules with different optical properties contained in the islet (e.g., lipids, fibers, and proteins), and to light scattering and/or absorption by the same molecules. To tackle this issue, ExM protocol variations^61,62^ that guarantee multiplexed localization, molecular specificity, and tissue clearing to increase light penetration^63^ and signal-to-noise ratio, potentially allowing rapid reconstruction of whole-stained islets, and hence perform a more accurate molecular interrogation in diabetic and non-diabetic tissues.

## Materials and Methods

### Cell culture

INS-1E cells (kindly provided by Prof. C. Wollheim, University of Geneva, Medical Center) were maintained in culture at 37°C, 5% CO_2_ in RPMI 1640 medium containing 11.1 mmol/L D-glucose, supplemented with 10% heat-inactivated fetal bovine serum (FBS), 10 mmol/L HEPES, 2 mmol/L L-Glutamine, 100 U/mL penicillin-streptomycin, 1 mmol/L sodium-pyruvate, 50 μmol/L tissue culture grade β-mercaptoethanol. For ExM experiments, cells were plated at 70% confluency on 18 mm coverglass and grown for 48 h. For lifetime experiments, cells were plated at 70% confluency onto sterilized and fluorescence-microscopy-suitable dishes (IbiTreat µ-Dish 35-mm, Ibidi) for 24-48 h. Then, cells were exposed to cytokines (IL-1β 10 U/ml and IFN-γ 100 U/ml diluted in the complete medium) for 24 h. For control samples, cells were washed and replaced with a fresh complete medium.

### Western blot

Western blot technique was used to quantify and compare the intracellular concentration of proteins in cytokine-treated INS1-E cells and in the untreated control. INS-1E cells were treated with Cytokines (IL- 1β 10 U/ml IFN-γ 100 U/ml diluted in 1 ml of medium RPMI) for 24 h. For western blotting, the cells were lysed in 95 °C Laemmli buffer (60 mM Tris-HCl pH 6.8, 2%SDS, 10% glycerol). Protein concentration was measured using Pierce™ BCA Protein Assay Kit (Thermo Fisher) following manufacturer’s instructions. Samples were then prepared by adding DTT (dithiothreitol) to a final concentration of 100 mM and bromophenol blue to the appropriate volume of cell lysate in order to load 20 ug of total lysate for each sample. Samples were subsequently subjected to SDS-PAGE and transferred onto a nitrocellulose membrane. Non-specific protein binding was blocked by incubation with 5% non-fat dry milk in PBST (phosphate buffer saline with 0,05% Tween-20) for 1 hour. Primary antibodies were incubated overnight at 4 °C in PBST with 5% milk with gentle shaking. After three washes with PBST, HRP-conjugated secondary antibodies were incubated for 1h at room temperature. Complete list of employed primary and secondary antibodies is reported in Table 1. After washing, membranes were incubated with ECL™ Prime Western Blotting System (GE Healthcare) and the chemiluminescence signal was detected using ChemiDoc MP Imaging System (BIORAD). Reported values from independent experiments are normalized on the GAPDH signal and presented as a percentage of the control sample (not treated).

### Live-cell imaging

For phasor-FLIM metabolic experiments, living INS-1E cells were examined after 24 h of cytokine exposure in the maintenance condition (RPMI 1640 medium containing 11.1 mmol/L D-glucose, supplemented with 10% heat-inactivated FBS, 10 mmol/L HEPES, 2 mmol/L L-Glutamine, 100 U/mL penicillin-streptomycin, 1 mmol/L sodium-pyruvate, 50 μmol/L β-mercaptoethanol) at 37 °C. For the glucose stimulation experiments, cells (control and cytokine-treated samples) were washed with SAB (114 mM NaCl, 4.7 mM KCl, 1.2 mM KH2PO4, 2.5 mM CaCl2, 1.16 mM MgSO4, and 20 mM HEPES (pH 7.4)) supplemented with 2.2 mM glucose, and then incubated with SAB 2.2 mM glucose for 45 min (low glucose concentration) at 37°C, 5% CO_2_. Then, glucose was added to reach a final concentration of 16.7 mM (high glucose concentration). The same cell clusters were acquired with 2-photon microscopy at low and high glucose concentrations. For iMSD experiments, INS-1E cells were treated with Cytokines (IL-1β 10 U/ml IFN-γ 100 U/ml diluted in 1 ml of medium RPMI) to investigate the diffusive motion of insulin granules after 24 hours of administration. Insulin granules were marked by ZIGIR, a fluorescent probe that labels Zinc-rich granules^41^. INS-1E cells were plated into 1 ml IBIDI plates (81156: μ-Dish 35 mm, high ibiTreat: Ø 35 mm, high wall) incubated at 37 °C and 5% CO_2_ for 24 hours. For the labeling process, cells were incubated 1 μM ZIGIR into the cells’ medium and incubated for 15 minutes.

### Cell fixation

Control and cytokine-treated cells were fixed according to the structure to stain. For tubulin, the specimens were incubated in the pre-extraction buffer consisting of 0.1 mM Pipes, 1 mM MgCl_2_, 1 mM EGTA, 0.5% Triton, and 3% paraformaldehyde (PFA) in dH_2_O for 1 minute at room temperature (RT) and then fixed with 3% PFA+0.1% glutaraldehyde (GA) for 10 min at RT, following three washes in 1x phosphate buffer saline (PBS), 5 min each. For the actin staining, the samples were fixed with 4% PFA in PBS for 30 min at RT and washed 3 times with PBS, 5 min each. For mitochondria, cells were fixed with 3% PFA+0.1% GA in PBS for 30 minutes at RT and washed 3 times in PBS, 5 min each. Finally, for the insulin granule staining, cells were fixed with 4% PFA in PBS for 30 minutes at RT and washed 3 times with PBS, 5 min each.

### Immunostaining and functionalization

After fixation, cells were permeabilized with PBS+0.1% triton X-100 (PBST) for 10 min at RT, washed 2 times with PBS, and then blocked with 2% bovine serum albumin (BSA) for 30-45 min at RT. The samples were incubated with the following primary antibodies: mouse anti-α-Tubulin (diluted 1:200 in PBST) for 2 h at RT; rabbit anti-Insulin and rabbit anti-TOM20 (diluted 1:200 and 1:100, respectively, in PBS+0.1% Tween) overnight at 4 °C. then, after 3 washes with PBST (for tubulin) or PBS+0.1% tween (for mitochondria and insulin) for 10 min each, the specimens were incubated with secondary antibodies anti-Rabbit Alexa Fluor 488 and anti-mouse Alexa Fluor 488, diluted 1:200 in PBST and PBS+0.1% tween, respectively (see Table 1). For Actin, we adapted the protocol developed by Park et al^64^. Shortly, cells were stained with Alexa Fluor 488-conjugated phalloidin (diluted 1:5 in PBST) for 1 h at RT. After washing three times in PBST, cells were stained for 60 min with an anti-fluorophore antibody (rabbit anti-Alexa Fluor 488 antibody, diluted 1:100 in PBST) for 1 hour at RT. Next, cells were washed three times with PBST. Cells were then stained for 2 hours with a secondary antibody (Anti-rabbit Alexa fluor 488, diluted 1:100 in PBST) (see Table 1). The stained samples for tubulin, actin, insulin, and mitochondria were then washed 3 times with PBS, 10 min each, and then functionalized with 0.1 mg/mL of 6- ((acryloyl)amino)hexanoic Acid, Succinimidyl Ester (AcX, ThermoFisher, A20770) for ≥ 6 h at RT. After the functionalization step, the samples were washed 3 times with PBS, 10 min each.

### Polymerization, digestion, and expansion

The stained and functionalized specimens were soaked in a gelled solution consisting of 8.625% sodium acrylate, 2.5% acrylamide, 0.15% N,N′-methylenebis(acrylamide), 2 M NaCl, 1× PBS, diH2O, 0.2% ammonium persulfate (APS) and 0.2% (v/v) tetramethylethylenediamine (TEMED), with TEMED and APS last added in this order. The gelation solution (~50 μL) is placed on the hydrophobic surface, and the coverglass is placed on top of the solution with cells facing down. Gelation is allowed to proceed at RT for 1-1.30 h. After gelation, the gels attached to the coverglass were removed and placed in a digestion buffer consisting of 50 mM Tris-HCl (pH 8), 1 mM EDTA, 0.5% Triton X-100, 1 M NaCl supplemented with 8 units mL^−1^ proteinase K added freshly. Gels were digested overnight at RT with gentle shaking and then washed with 1 μg/mL DAPI in PBS for 10 minutes. Then the hydrogel is moved into a 60 mm petri dish and soaked in ~50 mL DI water to expand it. Water is exchanged every 30 minutes and 4 times until expansion is complete. At the final expansion, small gel pieces were cut from the gelled samples and observed by confocal microscopy.

### Fluorescence microscopy

Unexpanded and expanded fixed samples stained for tubulin, actin, insulin, and mitochondria were acquired with an inverted Zeiss LSM 800 confocal microscope (Jena, Germany). The acquisition was performed by illuminating the sample with a 405, 488, and 561 nm laser using a 63×/NA 1.4 oil-immersion objective. DAPI, AlexaFluor 488, and AlexaFluor 568 fluorescence were collected between 410-510 nm, 510-590 nm, and 590-700 nm, respectively, with GaAsP detectors. Z-stacks with step sizes of 1 µm and 0.6 µm were acquired for insulin and tubulin staining, respectively, while tiled images were stitched into a mosaic (~895 µm x 895 µm) to find the same cell cluster in the unexpanded and expanded specimens. Stacks of images for iMSD experiments were performed with an inverted Zeiss LSM 800 confocal microscope (Jena, Germany) exciting the ZIGIR at 561 nm and collecting the fluorescence between 590 nm and 700 nm. The pinhole aperture was set at 1 Airy (53 μm) and a 63x/1.4 oil objective was used. 500 frames per image were collected and each frame was 256×256 pixels size, with acquisition time of 204.80 ms. Metabolic imaging was performed by an Olympus FVMPE-RS microscope coupled with a two-photon Ti:sapphire laser with 80-MHz repetition rate (MaiTai HP, SpectraPhysics) and FLIMbox system for lifetime acquisition (ISS, Urbana Champaign). NADH was excited at 740 nm and emission was collected using a 30×/NA 1.0 planApo silicon immersion objective in the 420-460 nm range. Flimbox system calibration was performed by measuring the characterized mono-exponential lifetime decay of Fluorescein at pH=11.0 (4.0 ns upon excitation at 740 nm, collection range: 480-570 nm). To prepare the calibration sample, a stock of 100 mmol/L Fluorescein solution in EtOH was prepared and diluted in NaOH at pH 11.0 for each calibration measurement. A 512×512 pixels image of FLIM data was collected into 25-30 frames, with an acquisition time typically of 1-2 minutes.

### Data analysis

For the expansion factor (EF) characterization, cells labeled for actin (for the cellular EF, n=15) and mitochondria (for subcellular EF, n=15) were acquired before and after expansion with confocal microscopy and calculated the EF by the ratio of cell and mitochondria area. The degree of sample deformation caused by ExM was calculated by comparing the pre- and post-Ex images stained for tubulin, actin, mitochondria and insulin (n=3 for each labeled structures) using the structural similarity index (SSIM index) plugin. For the count of insulin granules, Z-stack acquisitions with low downsampling along the Z-axis were performed to image different granules in each plane (Z step=1 µm). Then, Z-stacks were converted into a binary image by defining an appropriate threshold and analyzed with the “Analyze Particle” tool. The number of granules for a single cell was normalized for the cell area and then the percentage reduction of granules after cytokine treatment was calculated. For α-tubulin, Regions Of Interest (ROIs) of 10×10 µm^2^ at different planes along the Z-axis (basal and equatorial, see Results; n=26, from 3 independent samples) were selected and processed by Fiji to improve the signal-to-noise ratio (SNR) (Process˃Math˃Subtract Background). Then, the tracing process was performed by FiNTA, to generate binary images of the MT meshwork. Finally, Finta-processed images were converted by the Fiji Skeletonize plugin (Plugin˃Skeleton˃Skeletonize (2D/3D)) into a binary image and then analyze using “Analyze Skeleton”, to achieve information on the number of branches, junctions, triple points, quadruple points, endpoint, and average branch length. For the calculation of the MT opening (number and average area), the binary images were analyzed by MorphoLibJ plugin (Plugin˃MorphoLibJ˃Segmentation˃Morphological Segmentation˃Border Image). To provide a way of characterizing the intensity distribution of the MT network in terms of branches’ average distance and number of intensity peaks, an adaptation of Garlick’s method^48^ was implemented in Python 3.6. Such features were calculated through an image-processing algorithm plotting intensity values along 175 diameters running across the circle inscribed in every square image. Consequently, each sample was associated with an average number of intensity peaks and the average distance between peaks. To investigate the treatment response, the analysis was extended to an equal number (n = 76) of cytokine treatment and control experiments. For cortical actin, Finta was used to evaluate the number of branches, junctions, triple points, quadruple points, end points and the average branch length (then divided by the square root of the normalization factor, 2.17; n=28 cells). Due to the heterogeneity of the actin staining, the entire field of view, instead of 10×10 µm^2^ ROIs, was processed. Next, the data were divided by a normalization factor (calculated as follows: frame area/ expansion factor/ number of cells in the frame). MorphoLibJ plugin (see tubulin) was used to calculate the number and the size of the areas enclosed between branches (termed corrals). The number of corrals was then divided by the frame area (cell area/expansion factor). The average area of corrals was calculated and then divided by the expansion factor, 4.7. The average of the perimeters of corrals was calculated and divided by the square root of the expansion factor: 2.17. For the bleb analysis, an equal number of cells for control and the cytokine-treated samples (N=93) was analyzed through the observation of the actin-positive cytoplasmic protrusion. This value was then divided by the number of cells in the frame. The average number of blebs per cell was thus calculated. For the morphology analysis of mitochondria, regions with high-density mitochondria were selected, processed by Fiji to improve the SNR (Subtract Background and Gaussian Filter), and analyzed with the plugin MorphoLibJ (input “Object Image”), by setting an opportune tolerance value for each ROI. A binary image was produced and analyzed with the MorphoLibJ tool “Analyze”, obtaining information on the area, perimeter, and circularity of mitochondria. To obtain the dynamical and structural information (iMSD experiments), we employed a custom MATLAB script (MathWorks Inc., Natick, Ma) which computes the spatiotemporal correlation function by Fast Fourier methods. The script is fully described here in^40^. The iMSD curve derived from the script brings out the diffusion law and two diffusion parameters (α, Dm). These are key parameters to describe the diffusive motion: α for granules motion (α < 1 as anomalous diffusion, α = 1 as isotropic diffusion, α > 1 as guided diffusion) and Dm for the diffusivity inside confined regions. A more exhaustive description of the image processing can be found in^40^. For the phasor-FLIM analysis, metabolic maps were produced in Python 3.6 taking into consideration NADH-free and NADH-bound monoexponential characteristic fluorescence lifetime, to enable metabolic measurements in Phasor-FLIM. Indeed, our procedure is as according to [https://www.nature.com/articles/s41596-018-0026-5] with the following exception: letting the algorithm sample the phasor plot to automatically map metabolism ranges. In fact, the most extreme intersection points between the experimental point cloud and the straight line joining NADH-free and NADH-bound reference points can be determined geometrically. The next step is computing the distance between each data point and the line perpendicular to the metabolic segment passing through one of the two intersection points. The maps’ look-up tables come from linear segmented colormaps setting 10% and 90% values to anchor the colormap limits. Mann–Whitney test and Shapiro-Wilk normality test were performed for the statistical analysis. For the normally distributed data we choose the unpaired t test, instead for the data not normally distributed we choose the Kolmogorov-Smirnov test (nonparametric) (iMSD analysis). P-value<0.05 for statistical significance.

## Acknowledgments

This work has received funding from the European Research Council (ERC) under the European Union’s Horizon 2020 research and innovation programme (grant agreement No 866127, project CAPTUR3D). PM and MT receive support from the Innovative Medicines Initiative 2 Joint Undertaking under grant agreements No 115,797 (INNODIA) and 945,268 (INNODIA HARVEST). These Joint Undertakings receive support from the Union’s Horizon 2020 research and innovation program and “EFPIA”, “JDRF” and “The Leona M. and Harry B. Helmsley Charitable Trust”. We thank Fabio Azzarello for the assistance with Linux.

## Author contributions

L.A.P. performed experiments, analyzed data; V.D.L. and M.T. performed Western Blot experiments, analyzed data; M.B. performed cytoskeletal and phasor-FLIM analysis; S.G. performed iMSD analysis; P.M. designed research, wrote the manuscript; F.C. provided funds, designed and supervised research, wrote the manuscript; L.P. designed research, performed experiments, analyzed data, wrote the manuscript.

