## Supplementary Material for "Direct optical nanoscopy unveils signatures of cytokine-induced β-cell structural and functional stress"

### Title:

### Affiliations:



### Supplementary Figures

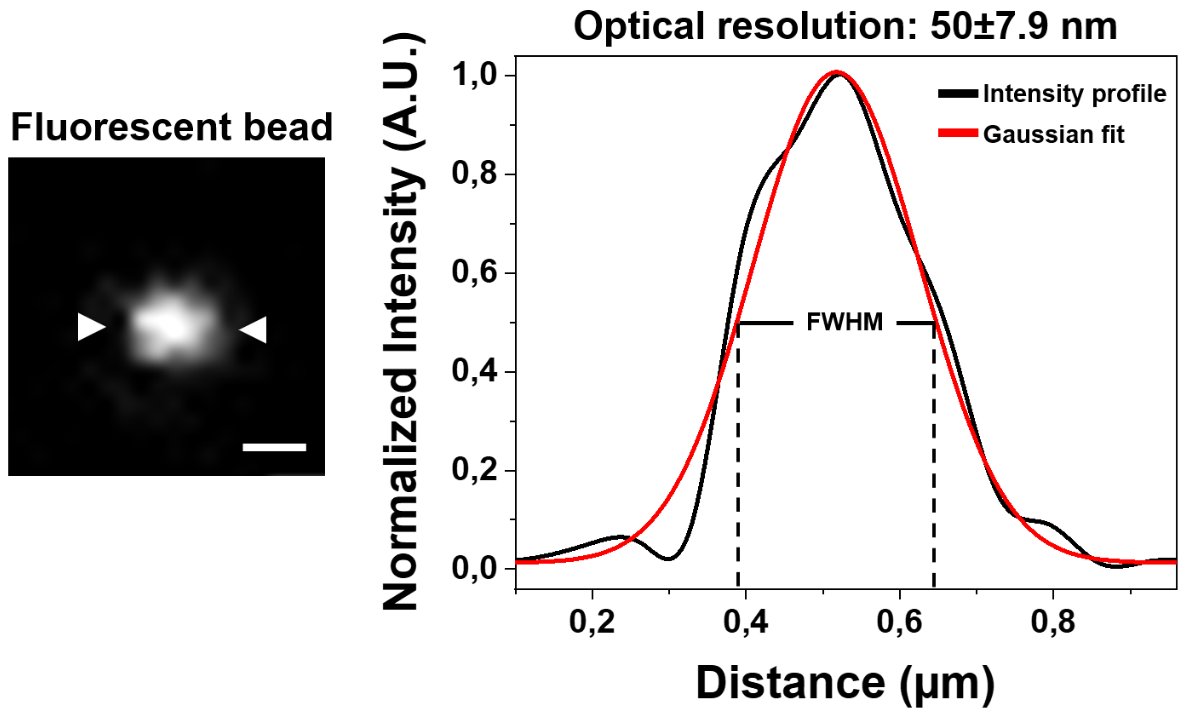


**Fig. S1. Optical resolution.** Confocal images of 100 nm fluorescent beads (TetraSpeck Microspheres, Invitrogen, T7279) to determine the optical resolution. The intensity profiles were fitted to a 2-D-Gaussian function to calculate the FWHM for individual beads shows (n. of beads= 3). Exc. 488 nm. Scale bar 0.3 µm.

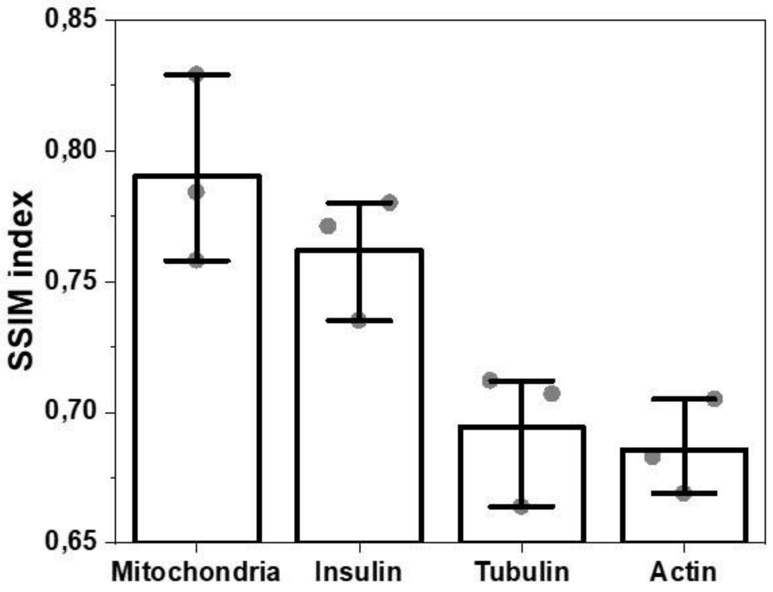


**Fig. S2. Structural similarity.** Comparison of the structural similarity between two confocal images using the structural similarity index measure (SSIM index; range value between -1 and 1; two identical images have an SSIM index close to 1). The data were shown as the mean ± SD (n=3 for each stained structure).

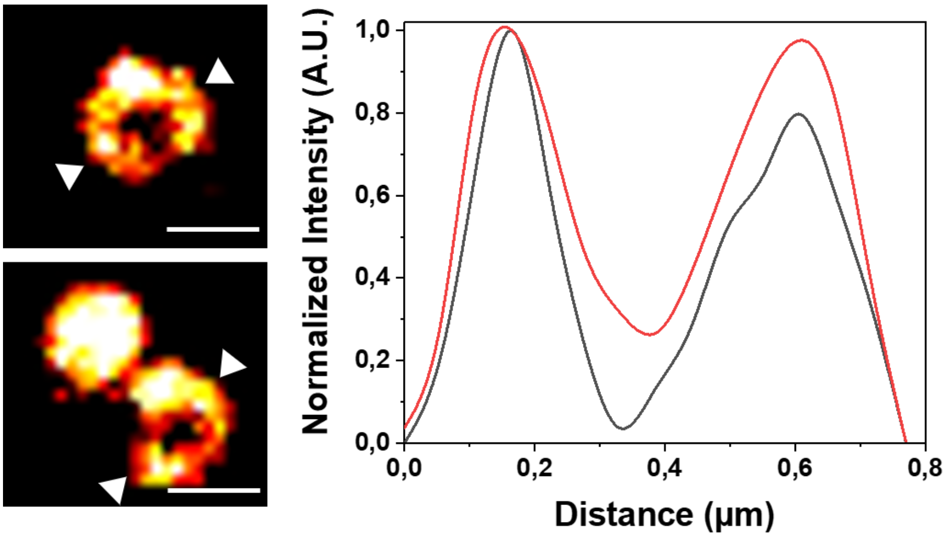


**Fig. S3. Insulin granule expansion.** Profile of the intensity of expanded insulin granules. The crystalline structure of insulin shows a doughnut-shape caused by the isotropic expansion of the labeled granules. Scale bar 0.5 µm.

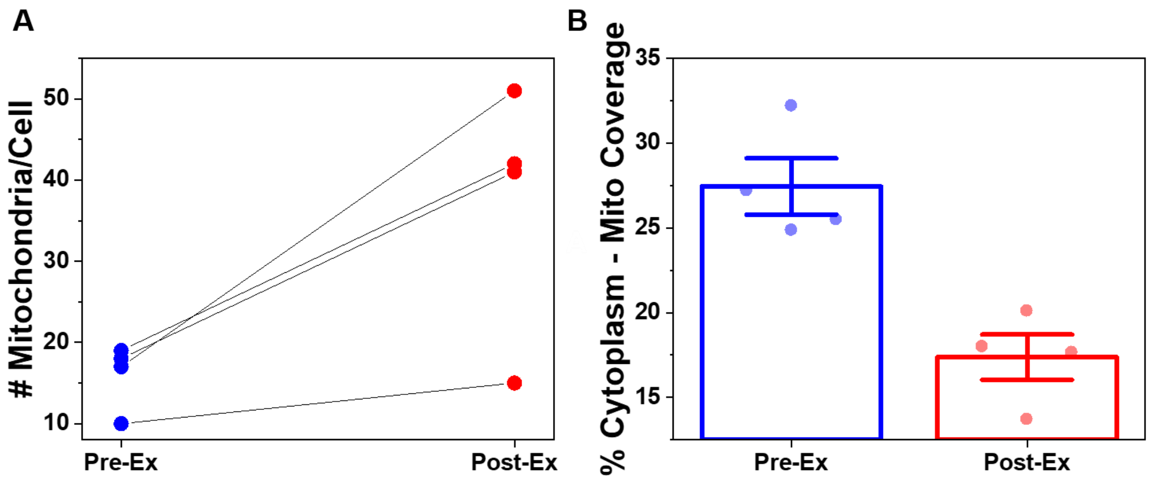


**Fig. S4. Mitochondria count and coverage analysis. A.** The number of mitochondria per cell in pre-(blue circle) and post-expansion samples (red circle) (N = 4 cells). **B.** The average percent cytoplasm area (cytoplasm area - nuclear area) which contains mitochondria.

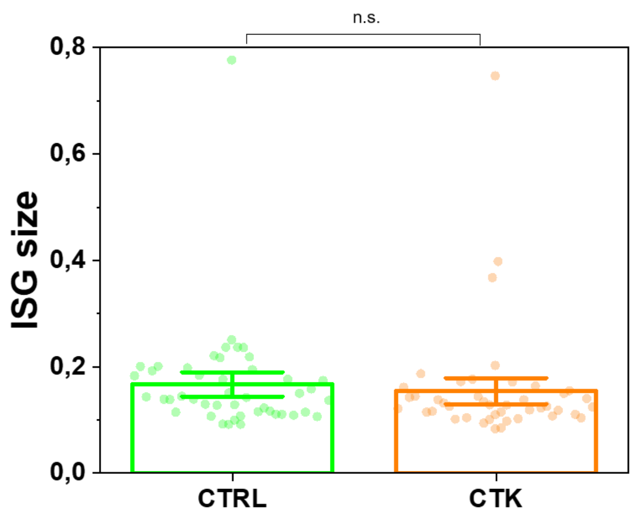


**Fig. S5. ISG apparent size in control and cytokine-treated samples.** Average size of ISG labeled with Zigir is not affected by cytokines treatment, shown as bar ± SE. A Mann–Whitney test was performed (n.s.= the two distributions are not significantly different).

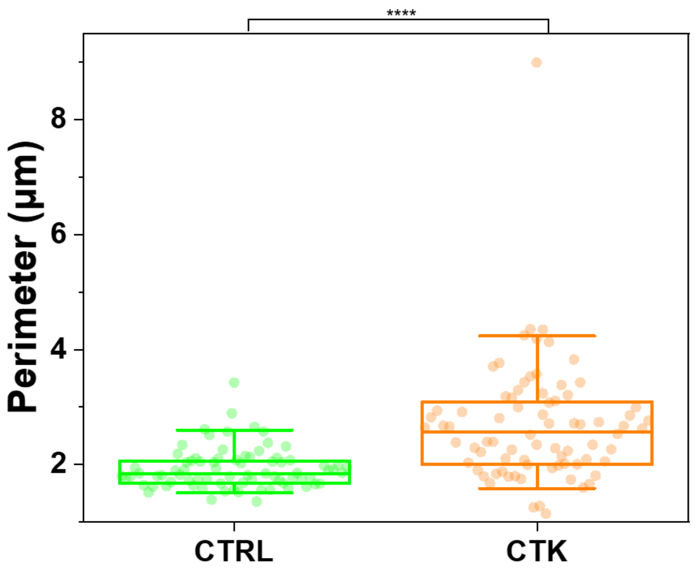


**Fig. S6. MT mesh perimeter.** Treatment with cytokine induces significant increase in corollas size, including mean perimeter. Data were presented as box plots with whiskers at the 5th and 95th percentiles, the central line at the 50th percentile, and the ends of the box at the 25th and 75th percentiles (n=78; 3 independent experiments).

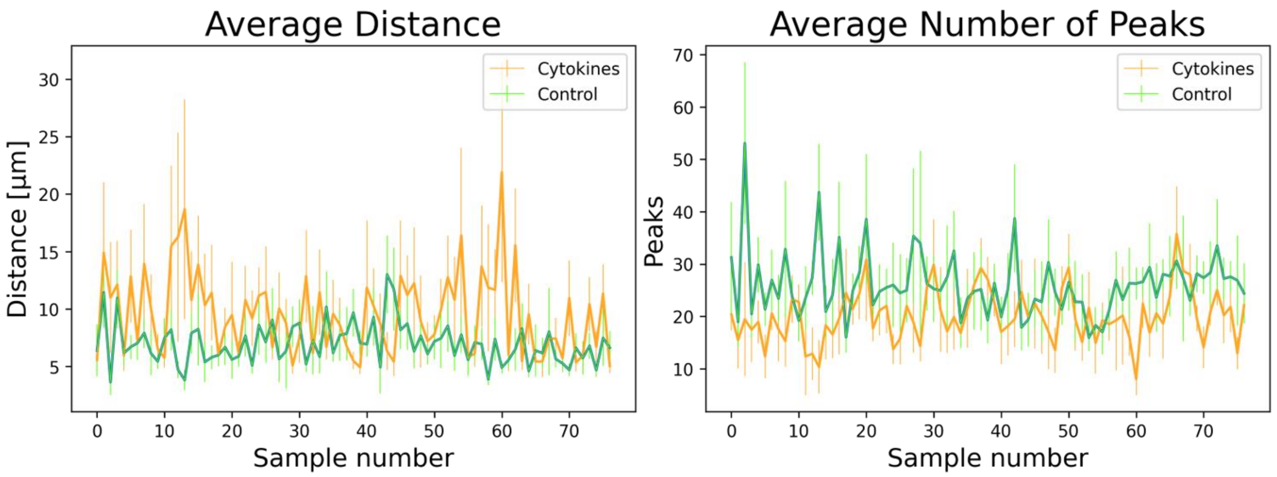


**Distance ratios:** 1.53 ± 0.19
**Peak ratios:** 0.75 ± 0.16

**Fig. S7. Intensity distribution analysis of MT mesh.** Example of the intensity plot profiles to evaluate the effect of cytokine treatment on each sample in terms of microtubule branching. The figure shows the average number of intensity peaks and the average distance between the peaks themselves, alongside standard deviation.

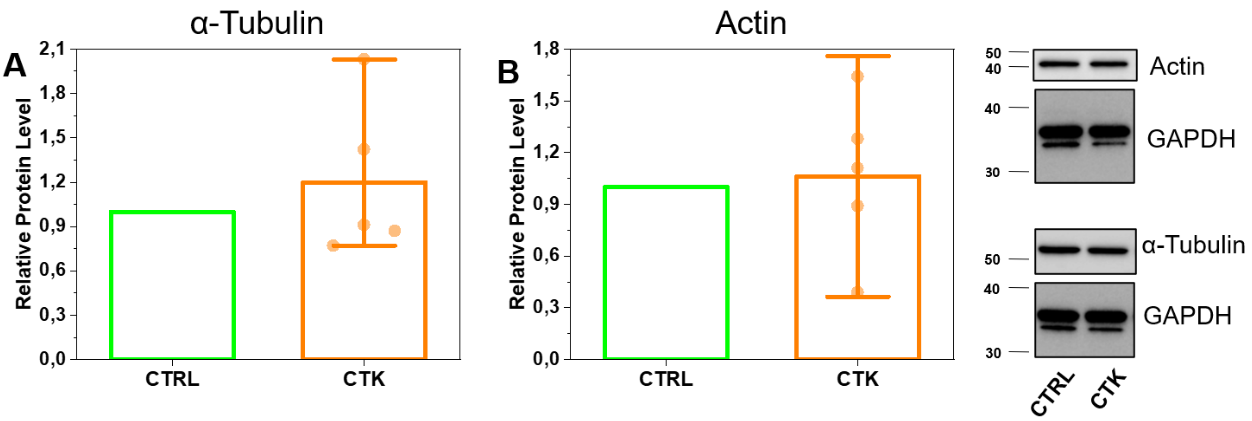


**Fig. S8. Western blot analysis.** Western blot analysis of tubulin (**A**) and actin (**B**). The differences were not statistically significant. n=5 independent experiments.

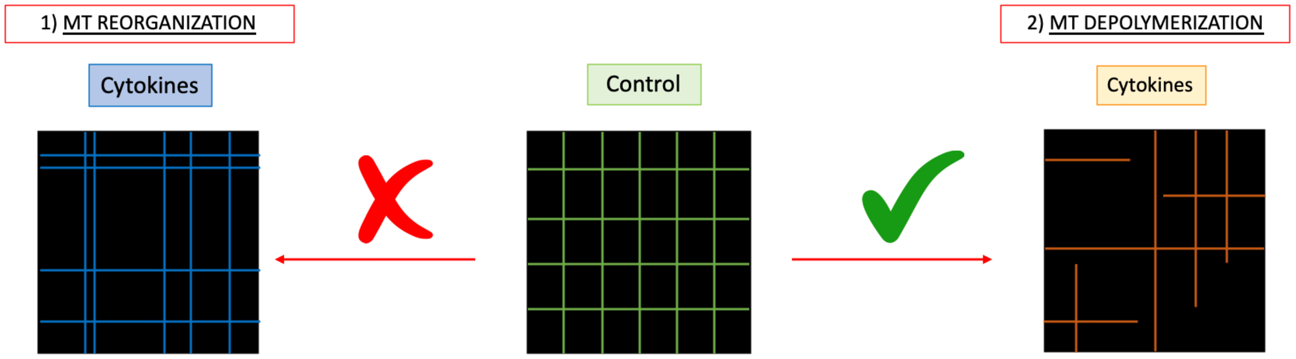


Table 1

| # Branches | | | # Junctions | | | # Quadruple Points | | | # Endpoint Voxels | | | Average Branch Length | | |
|---|---|---|---|---|---|---|---|---|---|---|---|---|---|---|
| 49 | 49 | 20 | 38 | 38 | 16 | 20 | 20 | 6 | 18 | 18 | 10 | 1,8 | 1,8 | 2,8 |

Table 2

| # Areas | | | Mean area | | | SEM area | | | Mean perimeter | | | SEM perimeter | | |
|---|---|---|---|---|---|---|---|---|---|---|---|---|---|---|
| 30 | 30 | 9 | 3,1 | 3,3 | 11,3 | 0,1 | 0,6 | 3,4 | 7,6 | 6,9 | 12 | 0,5 | 0,6 | 1,9 |

**Fig. S9. MT network modeling.** Schematic model of tubulin alterations after cytokine treatment. Data in table 1 were calculated manually. Data in table 2 were calculated with MorphoLibJ. These results demonstrate that the orange model is more affine to the ExM data.

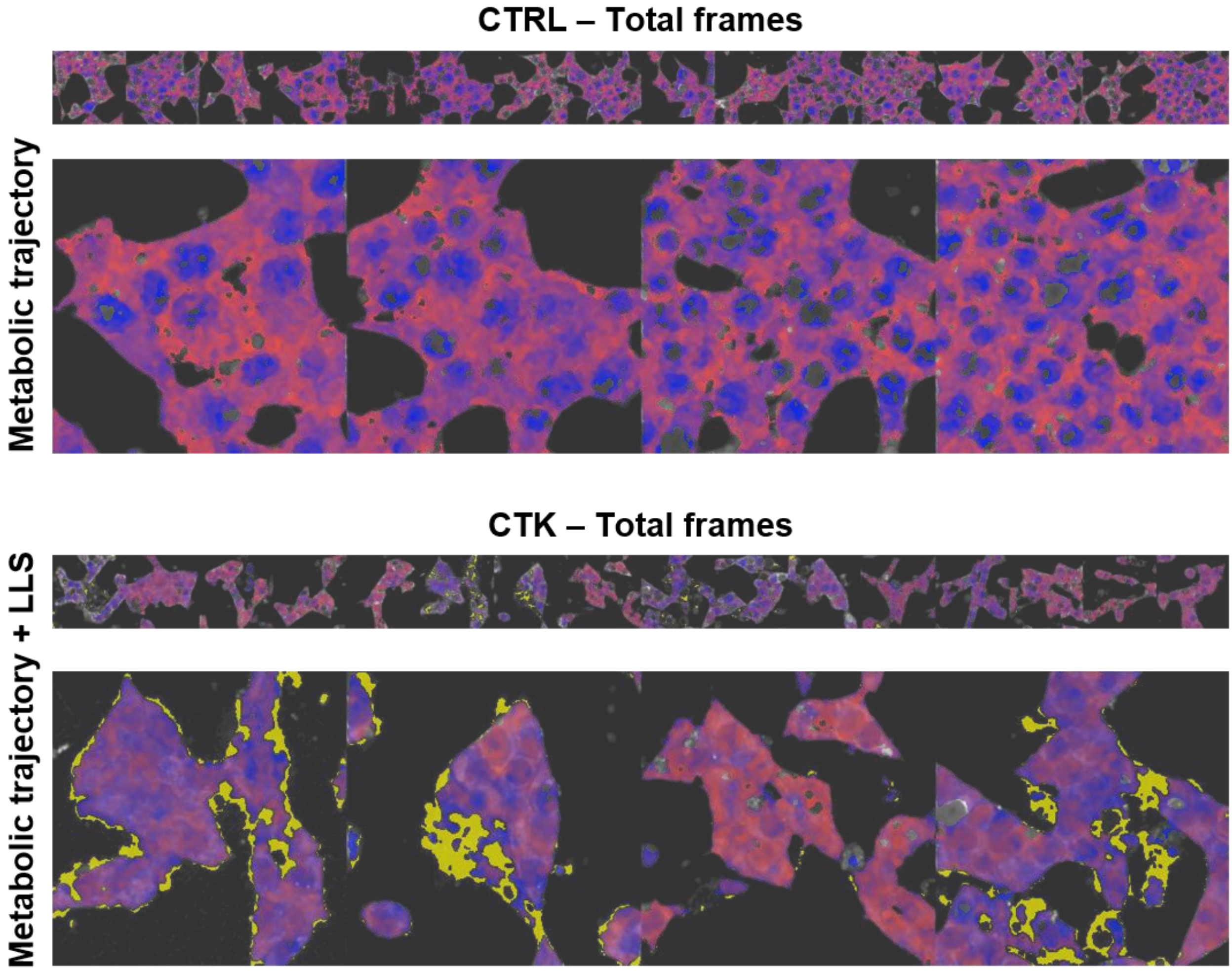


**Fig. S10. Phasor-FLIM signature of metabolic trajectory and oxidative stress.** Cytokine-treated samples show high pixel numbers characterized by long lifetime signature (yellow) respect with to the control sample.

| Molecule | Company | Cat. N. | Host | P/M | Dilution |
|---|---|---|---|---|---|
| **Anti-α-Tubulin Ab** | Sigma Aldrich | T6074 | Mouse | M | IF: 1:200 |
| **Anti-mouse IgG, AF 488** | Immunological Sciences | IS20014-1 | Donkey | P | IF: 1:100 |
| **Anti-mouse IgG, AF 568** | Immunological Sciences | IS20105-1 | Donkey | P | IF: 1:100 |
| **AF 488, Phalloidin** | Cell Signaling Technology | 8878S | | | IF: 1:5 |
| **Anti-Alexa Fluor 488 IgG** | Thermo Fisher Scientific | A-11094 | Rabbit | P | IF: 1:100 |
| **Anti-Rabbit IgG, AF 488** | Immunological Sciences | IS20015-1 | Donkey | P | IF: 1:100 |
| **Anti-Rabbit IgG, AF 568** | Immunological Sciences | IS20098-1 | Donkey | P | IF: 1:100 |
| **Anti Pig Insulin IgG** | BIO-RAD | 5330-0104G | Guinea Pig | P | IF: 1:200 |
| **Anti Insulin IgG** | Immunological Sciences | AB-84377 | Rabbit | P | IF: 1:200<br>WB: 1:1000 |
| **Anti-Guinea Pig IgG, AF 488** | Immunological Sciences | IS20017 | Goat | P | IF: 1:100 |
| **Anti-Guinea Pig IgG, AF 568** | Immunological Sciences | IS80102 | Goat | P | IF: 1:100 |
| **TOM 20 IgG** | Proteintech | 11802-1-AP | Rabbit | P | IF: 1:100 |
| **Tubulin alpha** | Immunological Sciences | AB-11661 | Rabbit | P | WB: 1:5000 |
| **Anti-Actin Antibody** | Sigma Aldrich | A 3853 | Mouse | M | WB: 1:3000 |
| **Goat Anti-Rabbit IgG** | BIO-RAD | 170-6515 | Goat | P | IF: 1:3000 |
| **Goat Anti-Mouse IgG** | BIO-RAD | 170-6516 | Goat | P | IF: 1:2500 |
| **Anti- GAPDH** | Sigma Aldrich | G8795 | Mouse | M | WB: 1:3000 |
| **DAPI** | Sigma Aldrich | D9542 | | | IF: 1:2000 |

**Table S1.** Antibodies and fluorescent dyes used for immunofluorescence (IF) and for Western Blot (WB), with their dilution.
